# Identification and prioritization of gene sets associated with schizophrenia risk by co-expression network analysis in human brain

**DOI:** 10.1101/286559

**Authors:** Eugenia Radulescu, Andrew E Jaffe, Richard E Straub, Qiang Chen, Joo Heon Shin, Thomas M Hyde, Joel E Kleinman, Daniel R Weinberger

**Author notes:** **Address correspondence to:** Daniel R. Weinberger, Lieber Institute for Brain Development, Johns Hopkins Medicine, 855 North Wolfe Street, Baltimore, Maryland 21205.

## Abstract

Schizophrenia polygenic risk is plausibly manifested by complex transcriptional dysregulation in the brain, involving networks of co-expressed and functionally related genes. The main purpose of this study was to identify and prioritize co-expressed gene sets in a hierarchical manner, based on the strength of the relationships with clinical diagnosis and with the polygenic risk for schizophrenia. Weighted Gene Co-expression Network Analysis (WGCNA) was applied to RNA-quality adjusted DLPFC RNA-Seq data from the LIBD Postmortem Human Brain Repository (90 controls, 74 schizophrenia; Caucasians) to construct co-expression networks and detect modules of co-expressed genes. After internal and external validation, modules of selected interest were tested for enrichment in biological ontologies, association with schizophrenia polygenic risk scores (PRS), with diagnosis and for enrichment in genes within the significant GWAS loci reported by the Psychiatric Genomic Consortium (PGC2). The association between schizophrenia genetic signals and modules of co-expression converged on one module showing a significant association with diagnosis, PRS and significant overlap with 36 PGC2 loci genes, deemed as tier 1 (strongest candidates for drug targets). Fifty-three PGC2 loci genes were in modules associated only with diagnosis (tier 2) and 59 in modules unrelated to diagnosis or PRS (tier 3). In conclusion, our study highlights complex relationships between gene co-expression networks in the brain and polygenic risk for SCZ and provides a strategy for using this information in selecting potentially targetable gene sets for therapeutic drug development.

## INTRODUCTION

Schizophrenia, the rubric for the most severe psychiatric syndromes in the psychoses spectrum, continues to be an enigmatic human condition. Large-scale genomic [1], transcriptomic [2] and epigenomic [3] studies have started to reveal not only the multifactorial biological substrate of schizophrenia (SCZ), but also new challenges especially from the clinical translational perspective. While it has long been clear that genes and their proteins do not act in isolation to build brain circuitries and maintain their functionality [4], the subtle and complicated genetic and environmental interactions that influence the development and function of the brain remain largely a mystery. Consequently, translating new discoveries into efficient therapies is probably the most difficult and frustrating endeavor in contemporary psychiatry.

Taking into account that transcriptional regulation plays a major role in neurodevelopment and neuronal activity [5], a promising approach to study genetic interactions and their implications in risk for schizophrenia is gene co-expression analysis. The logic behind this approach is that functional gene assemblies probably require a co-regulated transcriptional profile. Consequently, networks of co-expressed genes from postmortem brain gene expression data could mirror such functional gene assemblies. A popular bioinformatics tool for constructing and studying gene co-expression networks is Weighted Gene Co-Expression Analysis (WGCNA) [6]. This approach has been used to characterize patterns of co-expression in the normal brain [7], in autism spectrum disorders [8], in schizophrenia (human and animal modeling studies) [9–12] and across mental disorders [13]. Importantly, co-expression network analysis has also been used to understand the partitioning of polygenic SCZ risk in the brain transcriptome.

For example, in a recently published study, Fromer et al took a stepwise approach by combining transcriptomics and genetics techniques, including gene co-expression analysis, and identified a sub-network of co-expressed genes with roles in synaptic transmission that was highly enriched for SCZ genetic associations [14]. In another study, Pergola et al used a multi-modal approach including co-expression analysis and found that a co-expression profile including DRD2 and other SCZ risk genes was associated with intermediate phenotypes of schizophrenia [15]. While these earlier studies provide potentially important evidence for gene network associations in SCZ, they focused on rather particular aspects of genetic risk integration with co-expression in the brain transcriptome.

In the present study, we have taken a more global and stepwise approach to characterize brain networks of co-expressed genes and their association with the clinical state of schizophrenia and with polygenic risk of SCZ. We first perform a systematic characterization of gene co-expression in postmortem DLPFC tissue from controls (CTRL) and patients with schizophrenia (SCZ). We also critically address the potential influence of RNA quality on network association, which has not been specifically considered in earlier work. This is an important potential confounder as co-expression may be subsumed by co-degradation. Most importantly, we then develop a pipeline to identify the gradual convergence between the DLPFC co-transcriptome and SCZ genetic signals, in order to select and prioritize gene sets as potential drug targets in schizophrenia.

## MATERIALS AND METHODS

### General pipeline of data processing

#### Human Postmortem Tissue

We used postmortem human brain tissue from the LIBD Human Brain Repository (HBR) for testing and the CommonMind Consortium (CMC) brain collection for validation [16, 2]. Dorsolateral prefrontal cortex grey matter tissue from both collections was used for RNA extraction. The protocol of brain acquisition (location, legal authorizations, informed consent, clinical review/ diagnosis), pre-processing and tissue quality check is detailed in [16, 2]. Briefly, the samples selected included tissue from adults (age of death=16-80), healthy controls (CTRL; N=90; M/F=72/18) and patients with schizophrenia (SCZ; N=74; M/F=52/22) donors, Caucasians (CAUC), all with RIN≥7.

#### RNA-Seq data processing

For the LIBD dataset, RNA was extracted from DLPFC gray matter (BA9/46) and RNA sequencing libraries were constructed with the Illumina poly A+ kit; the resulting sequencing reads were aligned to the human genome (UCSC hg 19 build) with TopHat (v2.0.4) [17]; following alignment, the expression for genes and exons was summarized in counts based on Ensembl v75 [18], then converted to RPKM (Reads Per Kilobase of transcript per Million mapped reads) and normalized by log2+1 transformation. Normalized expression data from all samples were adjusted to remove unwanted variance potentially explained by RNA quality (i.e. technical or biological artifacts) (details below and in **supplementary material**). All analyses were performed on expression data quantified at the gene-level; consequently only genes with sufficient abundance (median RPKM ≥ 0.1 across all samples) were retained for analysis. This selection yielded 22,945 genes for the LIBD dataset and 27,779 genes for CMC data.

Processing CMC gene expression data is described in [2]. CMC library preparation utilized the Illumina Ribozero kit. We downloaded CMC BAM files from Synapse (https://www.synapse.org/); the BAM files were aligned with TopHat2 and expression of genes was quantified in counts relative to Ensembl v75 and subsequently converted to RPKM and normalized by log2+1 transformation.

#### Genotype data processing and Polygenic Risk Scores (PRS) calculation

DNA extracted from cerebellar tissue was processed and normalized as described in [16, 2]. Genotype imputation and quality check was performed with IMPUTE2 [19] and Shape-IT [20]. Only common SNPs in Hardy-Weinberg Equilibrium (at p>1e-6) with MAF>5% were retained for analysis [2].

Polygenic risk scores (PRSs) were calculated for each sample by summing the imputation probability of the reference allele of the clumped SNPs using PLINK v1.07 (http://pngu.mgh.harvard.edu/purcell/plink/) [21] and weighted by the natural log of the odds ratio from PGC2 GWAS results [1]. We used PRS based on 10 clinical SNP sets, corresponding to GWAS p values of p=5e-8 (PRS1), p=1e-6 (PRS2), p=1e-4 (PRS3), p=0.001 (PRS4), p=0.01 (PRS5), p=0.05 (PRS6), p=0.1 (PRS7), p=0.2 (PRS8), p=0.5 (PRS9), p=1 (PRS10) [22].

### Selection and prioritization of gene sets associated with risk for SCZ based on gene co-expression analysis

Processing the gene expression data to remove unwanted variability associated with sequencing and tissue confounders is presented in **figure 1-A1** and supplementary material.

**Figure 1:**
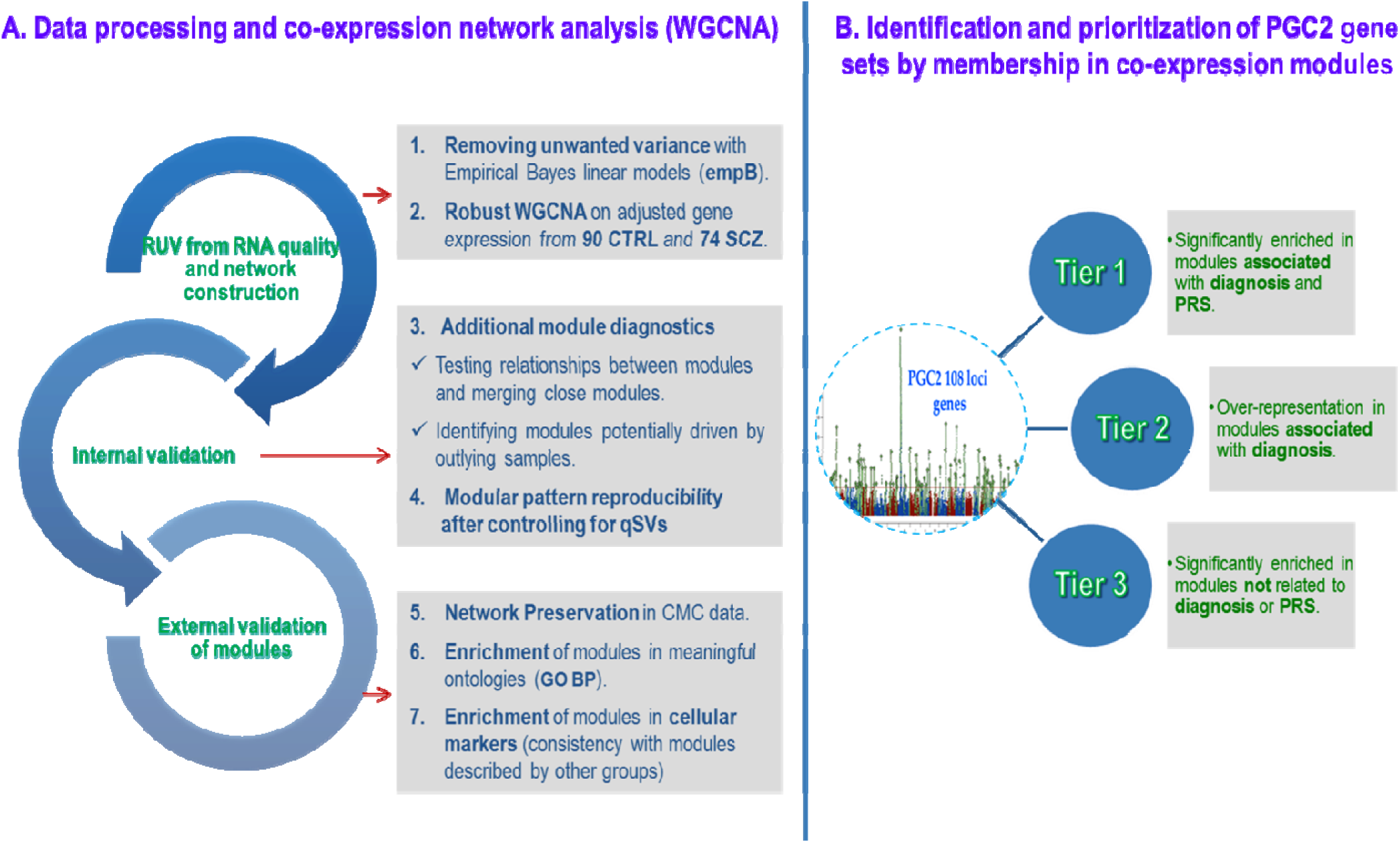
Analysis pipeline and criteria for selection and prioritization of gene sets associated with the genetic risk of schizophrenia. *A1*: Prior to co-expression network analysis, expression data normalized by (log2+1) transformation was adjusted to control for unwanted variation related to sequencing, tissue artifacts and population sub-structure (i.e. RIN, postmortem interval (PMI), exonic mapping rate, alignment rate and 10 genomic principal components-snpPCs) by using *empiricalBayesLM* function (WGCNA, version 1.61 (23)). *A2*: Co-expression network was constructed with WGCNA embedded routines (*blockwiseModules*) by applying the following parameters and procedures: bi-weight mid-correlation with “signed” networks to allow for potentially non-linear correlations between genes (24); β power=12 selected with “Soft Thresholding” function and applied to the gene correlations prior to network construction; modules of co-expression detection with hierarchical clustering using a measure of dissimilarity (the topological overlap); *A3*: Modules of co-expression were inspected through heatmaps of module specific gene expression across samples. In these heatmaps, well-defined modules are considered those displaying characteristic band structures, whereas the corresponding genes are highly correlated across samples; *A4*: Post-hoc WGCNA limited to 9239 genes organized in nine modules of interest after the primary network construction was performed. *A5*: External validation by using the *modulePreservation* function in WGCNA (23, 24), which computes network-based, pair-wise module preservation statistics by taking as input adjacency matrices in a reference set (the LIBD networks) and a test set (CMC); relevant statistics output: measures of preservation for density and connectivity summarized as individual composite Z scores (Z_summary_). By convention: Z_summary_ =0-2 means no preservation, Z_summary_ =2-10 means weak preservation and Z_summary_ >10 means strong preservation (27). *A6*: Enrichment in meaningful biological ontologies was tested with *enrichGO* function from the R package *clusterProfiler* (28): hypergeometric test was applied to test for over-representation of gene sets (i.e. module-specific genes) in relevant GO-BP; statistical threshold for significance was set at default values p=0.01 and q=0.05 with the Benjamini-Hochberg (BH) multiple comparisons correction method, using as background only the genes from the network construction, annotated by Entrez Gene IDs (which are largely protein-coding) (annotation performed with the Bioconductor package org.Hs.eg.db (29); *A7*: Enrichment in cellular markers was tested by using the hypergeometric test and gene lists embedded in the *userListEnrichment* function (24, 7, 30); *B*: Modules’ enrichment in PGC2 loci genes was also tested with the hypergeometric test (*userListEnrichment* function (24)); WGCNA intramodular analysis measures and functions were used to test associations between modules and polygenic scores and diagnosis (24).

The adjusted/”cleaned” expression data were then used as input for weighted gene co-expression analysis performed with functions implemented in the WGCNA package [23, 24].

Details about the next WGCNA steps are presented in **figure 1 (A2-7)** and **supplementary methods**. In brief:

#### 1. The co-expression network

based on empB-adjusted expression data was created for the combined sample of CTRL+SCZ with WGCNA. This method uses correlation between pairs of genes to construct co-expression modules. The modules can then be summarized by the first principal component (i.e. the “eigengene”) for each module (ME) [7, 23, 24]. The MEs can be regarded as expression profiles that best characterize the gene correlations within modules. Biological inference can be drawn from the genes in the constructed modules by using gene set enrichment analyses and by correlating module eigengenes with biological covariates. Likewise, intramodular analysis can be used to assess the degree of connectivity for the genes within modules and the gene-wise significance relative to association with traits of interest or diagnosis. The major advantage of MEs’ is in dimensionality data reduction, which makes them particularly suitable for correlation with traits of interest by eliminating the problem of multiple comparison corrections.

#### 2. Internal validation

a. Additional diagnostics of the co-expression modules was also performed (**figure 1-A3**). Previous studies in our group have highlighted the importance of RNA quality in gene expression analysis, especially in detection of differential diagnosis effects [25]. However, selecting the best method for controlling for these effects is not straightforward and relies much on the type of analysis performed. Methods using empirical Bayes moderated regression (e.g. ComBat [26] or *empiricalBayesLM* function used by us in this study) are stringent but they perform correction based only on “known” sources of unwanted variance. One possibility to remove variability from “unknown” sources of technical error can be performed by modelling “unknown” (latent) variables with “quality” surrogate variables (qSVs) [25]. This approach is based on data from a human brain RNA degradation experiment. Here, in addition to our main network construction with “cleaned” data after removing unwanted variance for “known” sources, we performed a complementary post-hoc analysis (**figure 1-A4**) with the purpose of confirming the modular pattern of the main network after adjusting also for unknown sources of variance represented by qSVs calculated from the “degradation matrix” [25] of the entire sample (90 CTRL + 74 SCZ).

#### 3. External validation

a. Module preservation analysis was performed in the CMC expression data selected and treated similarly with LIBD data (same parameters for RIN, age and application of empB adjusting for the unwanted variation) (**figure 1-A5**). This approach strengthened the comparability between the two datasets, necessary for testing the preservation. Moreover, module preservation in the context of the different library protocols in the LIBD and CMC datasets further supports their validity (details in **supplementary methods** and [27].
b. Further external validation was represented by enrichment in putatively meaningful ontologies and comparisons with modules previously identified by other groups. We tested for enrichment in Gene Ontology-biological processes (GO-BP) with functions implemented in *clusterProfiler* R package [28] (**figure 1-A6**).
c. As a further approach to module validation, we sought to see if our modules are similar to those reported in the study by Oldham et al (2008), which provided the initial characterization of the gene co-expression relationships in the human brain [7]. For this analysis, we tested for enrichment of our modules in cellular markers based on the cortex modules reported in [7] and additionally, on a transcriptome database including markers for neurons, astrocytes and oligodendrocytes by Cahoy et al (2008) [30] (**figure 1-A7**).

#### 4. Identification of gene sets associated with the genetic risk for SCZ within the co-expression networks

**Figure 1-B** describes a hierarchical approach of grouping gene sets within the co-expression modules by association with genetic risk for SCZ informed by the Psychiatric Genomic Consortium Genome Wide Association Study (PGC2 GWAS) [1].

We consider three tiers of gene sets relevant for association with SCZ biology, determined by the gradual convergence of biologically relevant function, clinical state and genetic risk. All the genes in the three tiers are PGC2 protein-coding genes within the 108 loci with GWAS significant genetic signal for risk of SCZ. **Tier 1**, i.e. the highest priority, comprises PGC2 loci gene sets that are enriched in modules significant for the association with both diagnosis and polygenic risk scores. **Tier 2** includes genes enriched in modules associated with the diagnosis but not with PRS. Finally, **tier 3** includes only the PGC2 genes over-represented in any module that was not associated with diagnosis or with the PRS.

We calculated the enrichment of PGC2 protein-coding genes (obtained from supplementary table 2 in [1]) in the entire co-expression network (**figure 1-B**). Two groups of genes were compared: one represented by all protein-coding genes used for the network construction annotated by gene symbol (N=15,359/ 22,945) and labeled by the color of their corresponding module and one represented by 309/349 PGC2 GWAS significant loci protein-coding genes according to the supplementary table 2 from [1]. We selected only 309 protein-coding genes because 40 had especially low abundance in our data set (RPKM<0.1) and therefore were not analyzed. For every pair of lists overlaps tested, the output was represented by uncorrected and FWE corrected p-values and by overlapping genes, respectively the PGC2 GWAS significant loci protein-coding genes represented in a module.

**Table 1:** Summary of “Gene Ontology Biological Processes” (GO-BP) and cell markers enriched in modules of co-expression.

| Modules and top 10 GO-BP enrichments |  |
| --- | --- |
| **Black- Astrocytes** |  |
| sensory organ development | astrocyte differentiation |
| eye development | carboxylic acid catabolic process |
| small molecule catabolic process | camera-type eye development |
| gliogenesis | organic acid catabolic process |
| glial cell differentiation | renal system development |
| **Blue- Neuron 1** |  |
| neurotransmitter secretion | vesicle-mediated transport in synapse |
| signal release from synapse | regulation of neurotransmitter levels |
| presynaptic process involved in chemical synaptic transmission | regulation of neuron projection development |
| modulation of synaptic transmission | neurotransmitter transport |
| neuron projection morphogenesis | synaptic vesicle transport |
| **Brown- Neuron 2** |  |
| single-organism behavior | learning |
| learning or memory | potassium ion transport |
| cognition | regulation of ion transmembrane transport |
| behavior | regulation of membrane potential |
| homophilic cell adhesion via plasma membrane adhesion molecules | modulation of synaptic transmission |
| **Green- Mitochondria 1** |  |
| protein targeting to ER | ribonucleoprotein complex biogenesis |
| translational initiation | protein localization to endoplasmic reticulum |
| establishment of protein localization to endoplasmic reticulum | mitochondrial ATP synthesis coupled electron transport |
| SRP-dependent cotranslational protein targeting to membrane | ATP synthesis coupled electron transport |
| cotranslational protein targeting to membrane | oxidative phosphorylation |

**Table 1 (cont.): Summary of “Gene Ontology Biological Processes” (GO-BP) and cell markers enriched in modules of co-expression**
| Modules and top 10 GO-BP enrichments |  |
| --- | --- |
| **Magenta- Mitochondria 2** |  |
| nuclear-transcribed mRNA catabolic process, nonsense-mediated decay | establishment of protein localization to endoplasmic reticulum |
| SRP-dependent cotranslational protein targeting to membrane | mRNA catabolic process |
| cotranslational protein targeting to membrane | protein localization to endoplasmic reticulum |
| nuclear-transcribed mRNA catabolic process | viral transcription |
| protein targeting to ER | viral gene expression |
| **Red- Oligodendrocytes** |  |
| ensheathment of neurons | negative regulation of neurogenesis |
| axon ensheathment | negative regulation of cell development |
| myelination | gliogenesis |
| oligodendrocyte differentiation | negative regulation of nervous system development |
| glial cell differentiation | negative regulation of cell differentiation |
| **Salmon- Microglia** |  |
| lymphocyte activation | immune response-activating signal transduction |
| positive regulation of immune response | adaptive immune response |
| activation of immune response | regulation of leukocyte activation |
| immune effector process | regulation of cell activation |
| immune response-regulating signaling pathway | leukocyte cell-cell adhesion |

**Table 2:** Summary of the relationships between co-expression modules and diagnosis, PRS and intra-modular PGC2 loci genes distribution.

| Modules | Association with dx | Association with PRS | Enrichment in PGC2 loci genes |
| --- | --- | --- | --- |
| Black | yes | yes | no |
| Blue<sup>3</sup> | no | no | yes |
| Brown<sup>1</sup> | yes | yes | yes |
| Cyan** | yes | yes | no |
| Green<sup>3</sup> | no | no | yes |
| Greenyellow | no | no | no |
| Magenta | no | no | no |
| Midnightblue* | yes | yes | no |
| Red | ± | no | $p_{\text{uncorr}} < 0.05$ |
| Salmon | ± | no | no |
| Turquoise<sup>2</sup> | yes | ± | $p_{\text{uncorr}} < 0.05$ |
| Yellow<sup>2</sup> | yes | no | $p_{\text{uncorr}} < 0.05$ |

*Legend:* <sup>1</sup>Module used for tier 1 genes selection; <sup>2</sup>Modules used for tier 2 genes selection; <sup>3</sup>Modules used for tier 3 genes selection; \*small-sized module with no GO-BP enrichment and weakly preserved in CMC data; \*\*module with no GO-BP enrichment.

#### Prioritizing the PGC2 loci genes distributed in the modules of co-expression

To prioritize the PGC2 protein-coding genes enriched in our co-expression modules, we used intramodular analysis routines from the WGCNA package [23, 24]. We first defined a measure of “Gene Significance” (GS) represented by the absolute value of Pearson correlation between each gene’s expression and diagnosis status. By averaging the GS for each module, we obtained a measure of “Module Significance” (MS). A module with high significance for diagnosis would be a module with many genes strongly correlated with the diagnosis. We then plotted the measure of module significance to visualize the most relevant modules for the association with the diagnosis (modules above a conventional cut-off=0.15 were considered significant [23, 24].

We next identified modules whose eigengenes (MEs) were significantly correlated with the SCZ PRS calculated for various P value thresholds as previously specified. We focused on modules correlated especially but not exclusively with PRS5-PRS6 because these scores presumably contain most of the true positive risk PGC2 genes and explain the maximum diagnostic liability in the PGC sample. After evaluating the relationship between MEs and PRS and diagnosis, we looked if the PGC2 protein-coding genes enrichment was significant in any of these modules, in order to select the overlapping PGC2 genes and prioritize them according to our criteria presented in **figure 1-B**.

## RESULTS

After expression data pre-processing, RNA-quality correction and network construction, 12 gene co-expression modules were identified in the overall sample with size between 40 and 1813 genes (N=12,475, 54%, were “grey” genes, not assigned to a module). The module specific gene distribution is reported in **supplementary table 1.**

The post-hoc WGCNA performed on the expression of selected genes after controlling also for hidden quality surrogate variables (qSVs) as described in **supplementary material**, yielded also a pattern of 12 modules, seven of which were enriched for ontologies and cellular markers similar with the “core” modules identified in the primary co-expression network analysis (**supplementary table 2**). In general, we found a significant degree of overlap between the modules of interest defined from the entire gene set network and the post-hoc qSVA constructed modules based on the selected genes (**supplementary table 2** and **supplementary figure 1**). We note, however, that qSV correction, a well validated method for differential expression and eQTL analysis [25], has not been critically examined in the context of network analysis, and this conservative approach results in a higher proportion of genes unassigned to modules (“grey” genes) (unpublished observations).

Additional results of modules diagnostics for the primary co-expression network are also reported in the **supplementary figures 2-5**, which show for the majority of modules (except cyan) characteristic band structures suggestive for well-defined modules, consistent across samples as described previously [23, 26]. Preservation analysis performed for the primary network constructed from 22,945 genes indicated that all 12 modules were preserved in the CMC data (**supplementary figure 6**). The preservation statistics varied between Zsummary=10 (“midnightblue” module) and Zsummary=60 (“yellow” module).

**Figure 2:**
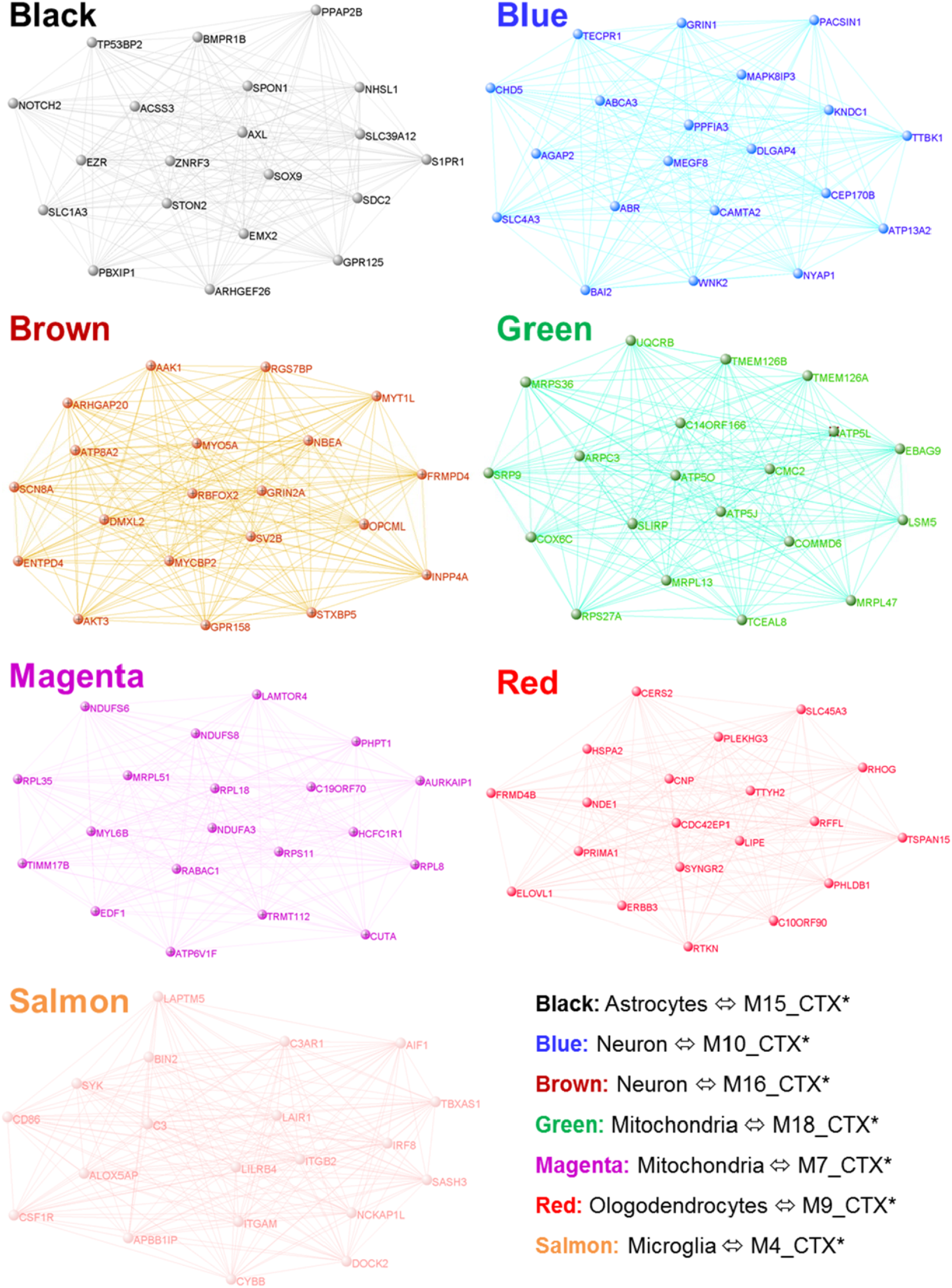
Top twenty most connected genes in seven “core” modules significantly enriched for biological processes relevant for cortex structure and functionality. Modules are represented by their colors assigned through co-expression network construction and module detection. *Legend:* * modules reported by Oldham et al (2008); modules’ representation created with VisAnt (http://visant.bu.edu/).

### General characterization of modules in DLFPC: enrichment in GO-BP and cellular markers

**Figure 2** and **table 2** summarize the significant and biologically plausible modules of gene co-expression identified in our networks based on CTRL and SCZ DLPFC RNA-Seq data. In general, these modules are not diagnostically specific and presumably reflect patterns of gene co-expression in human adult dorsolateral prefrontal cortex grey matter. Modules reflecting similar biological processes have been observed in earlier work in human brain [7]. Notably, we found seven “core” modules reproducing fundamental processes for nervous system development (i.e., neuronal differentiation and migration, synaptogenesis, gliogenesis and myelination) and functionality, metabolic processes critical for cellular survival and function, including nervous cells, specific neuronal processes linked to neuronal excitability and synaptic activity, immune system functions, mechanisms of transcription and translation, etc. A complete list with GO-BP enrichment for the empB modules is presented in **supplementary table 2**. Importantly, the organization of our co-expression modules significantly overlapped with the cell type specific modules reported in cortex by Oldham et al [7] (**figure 2, table 2, supplementary table 3**).

From our empB modules not enriched for GO-BP, but significantly enriched for cellular markers, we note: turquoise (overlapping with ‘neuron-M16_CTX’, M18_CTX, M11_CTX, ‘interneurons-M17_CTX’), yellow (overlapping with ‘glutamatergic synaptic function-M10, M18_CTX, M19_CTX), and greenyellow (‘olygodendrocytes-M9_CTX’, ‘glutamatergic synaptic function-M10, and ‘astrocytes-M15_CTX’). We also note that enrichment in cellular markers for these modules was more mixed than for the seven “core” modules (**supplementary table 3**).

### Identification of gene sets associated with genetic risk (PRS) for SCZ within the co-expression networks

Overall, we found that almost half of GWAS significant PGC2 loci protein-coding genes (148/309) were distributed in modules across the co-expression networks; 36 of them (which we designate as **tier 1 genes**) were in modules related to both diagnosis and polygenic risk score, 53 of them were in modules related only to diagnosis (**tier 2**) and 59 were in modules unrelated to diagnosis or PRS (**figure 3C**).

**Figure 3:**
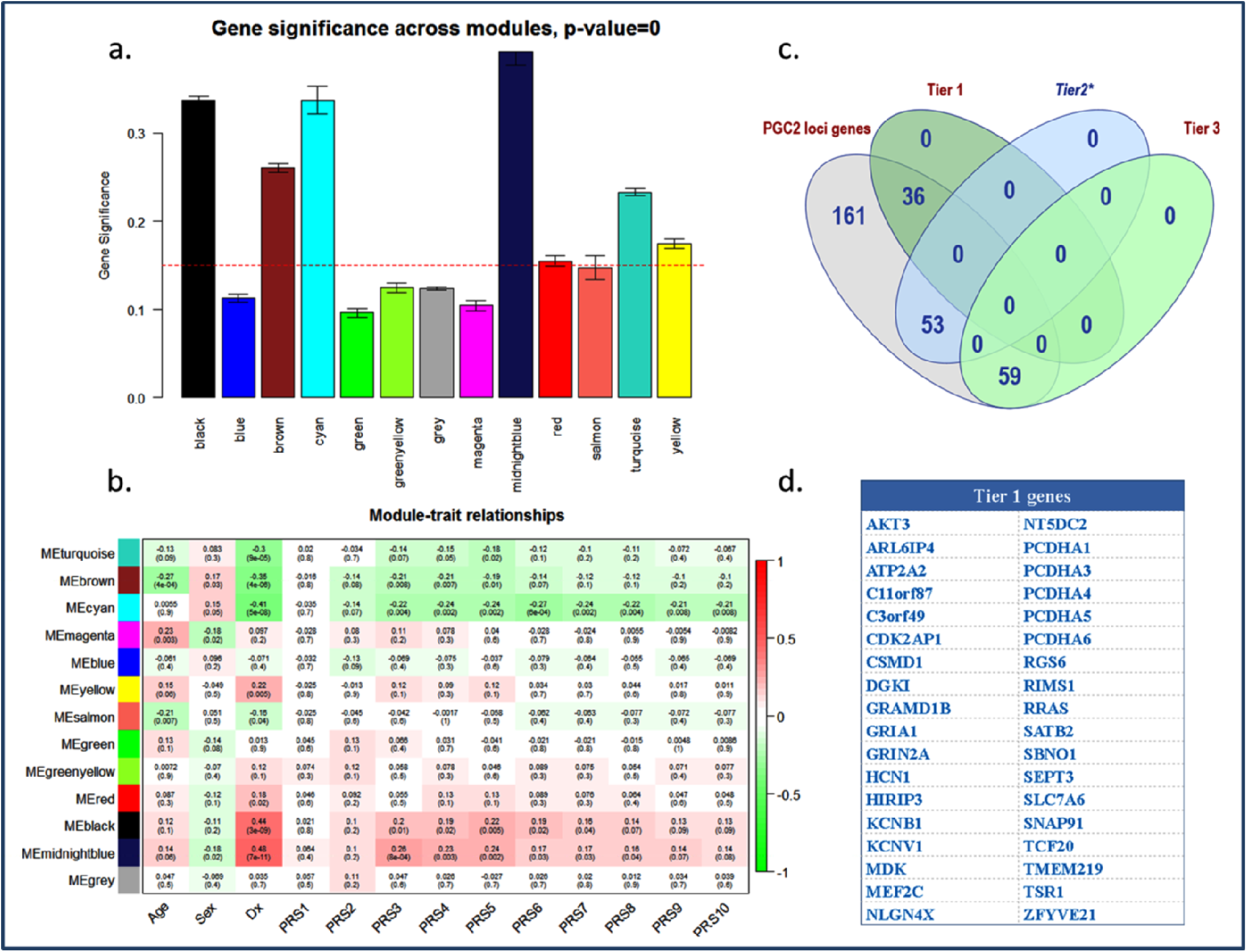
Prioritization of PGC2 loci genes based on the distribution in modules associated with polygenic risk score and diagnosis of SCZ. *a.* Modules significantly correlated with the diagnosis of SCZ; *b.* Modules’ eigengenes (MEs) correlated with diagnosis and PRS; *c.* Partitioning of PGC2 loci genes in tiers 1-3; *d.* Tier 1 genes*. Legend:* * these genes were members of two modules that shortly failed the significance for PGC2 loci genes enrichment after FWE multiple comparisons correction.

We found that MEs of four modules were significantly correlated with both PRS sets and diagnosis: the black module (enriched for astrocytes markers), the midnightblue module (mixed cellular markers), the brown (enriched for neuronal markers) and the cyan module (mixed cellular markers) (**figure 3B**).

Modules correlated only with diagnosis were: the yellow module (enriched for markers of glutamatergic synaptic function) and marginally, the salmon module (enriched for microglial markers) and the red module (enriched for oligodendrocytes markers) (**figure 3A**). Modules with a significant over-representation of PGC2 loci genes were blue, brown, green (p corrected values <0.05) and red, turquoise, yellow (p uncorrected values <0.05) (**table 2**). A synthesis of the associations between modules and diagnosis and polygenic scores, combined with the information regarding the PGC2 loci genes distribution within modules is presented in **table 2**.

### Tier 1 genes: PGC2 loci genes over-represented in modules associated with both diagnosis and PRS

We selected as **tier 1** 36 PGC2 loci protein-coding genes that were represented in the brown module (**figure 3d**) because this module’s eigengene is associated both with diagnosis and with PRS and brown is the only module with ME association with both of these features and also with PGC2 locus gene overrepresentation. Notable tier 1 genes are AKT3, essential in brain development [31], ATP2A2 (encoding for a magnesium-dependent enzyme that catalyzes ATP’s hydrolysis [32], genes involved in neural circuits development via cell adhesion processes (protocadherins alpha cluster-PCDHA1,3,4,5,6, ZFYVE21, RRAS), transcription factors (TCF20, MEF2C), regulators of G-protein signaling (RGS6), RRAS-a small GTPase involved in cell adhesion and axon guidance, and TSR1 (Ribosome Maturation Factor), which is important for fundamental functions related to protein synthesis and gene expression in all cells [32]. We also note among tier 1 genes two glutamatergic ionotropic receptors (GRIA1 and GRIN2A), and several potassium ion channels receptors (HCN1-Hyperpolarization Activated Cyclic Nucleotide Gated Potassium Channel 1, KCNB1-Potassium Voltage-Gated Channel Subfamily B Member 1 associated with epilepsy, and KCNV1-Potassium Voltage-Gated Channel Modifier Subfamily V Member 1, essentially expressed in the brain) [32].

### Tier 2 genes: PGC2 loci genes over-represented in modules significant for the diagnosis of SCZ but not PRS

We selected 53 PGC2 genes as tier 2 genes, with a trend of over-representation in turquoise and yellow modules (**supplementary table 4**). However, we specify that PGC2 enrichment did not reach the levels of significance after FWE correction for multiple comparisons (uncorrected values were p=0.023 for yellow, respectively p=0.01 for turquoise (**supplementary table 4**). As previously mentioned these modules represented mixtures of cellular markers (**supplementary table 3**) and were not significantly enriched for GO-BP. However, we considered them interesting modules based on the PGC2 genes present in these modules. One of the notable tier 2 genes in this module is ZNF804A [33]. Other interesting members of tier 2 genes are NDUFA13, NDUFA4L2, NOSIP, and NRGN.

### Tier 3 genes: PGC2 genes over-represented across networks

59 PGC2 GWAS significant loci genes belonged to **tier 3**: they were over-represented in blue and green modules that were neither associated with PRS, nor with the diagnosis. Of note, blue is one of the neuronal modules and green is enriched for mitochondria markers (**figure 2**). Interesting tier 3 genes are CACNA1C, CHRM4, FURIN, TSNARE1 (possibly implicated in neurotransmitter release by regulating the SNARE-Synaptotagmin complex), FXR1 (FMR1 Autosomal homolog, associated with Fragile X syndrome) [32]. The complete list of the tier 1-3 PGC2 loci genes is presented in **supplementary table 5**.

Finally, 161/ 309 PGC2 locus genes were randomly distributed in modules that did not pass the threshold of significance for PGC2 enrichment or were “grey” genes (not assigned to a module in any network). A notable “grey” PGC2 locus gene was DRD2, the best established drug target for SCZ. While not excluding the possible implication of any PGC2 gene in the etiopathogenesis of SCZ, based on our results, we believe that **tier 1** and **tier 2** genes, because of their association at least with illness diagnosis, are more attractive candidates for experimental studies to decipher pathophysiological mechanisms of SCZ or drug development.

## DISCUSSION

We have performed an extensive analysis of gene co-expression architecture in adult postmortem DLPFC from CAUC unaffected donors (CTRL) and from donors diagnosed with schizophrenia (SCZ). The main purpose of the study was to identify and prioritize co-expressed gene sets in a hierarchical manner, based on the strength of the relationships with clinical diagnosis and with the polygenic risk for schizophrenia. For this purpose we focused on co-expression modules that included PGC2 protein-coding genes, i.e. the genes within the significant 108 loci reported in the latest published GWAS of SCZ [1]. The rationale of our approach was that finding a convergence between the co-expression architecture in a region with known abnormalities in SCZ (i.e. DLPFC) with both the illness state and with genomic risk for the illness is a more optimal strategy to isolate potentially co-functional gene sets that could be investigated as harboring novel drug targets in SCZ. Importantly, we found seven “core” modules enriched for meaningful ontologies and significantly overlapping with modules reported by other groups. We further selected and hierarchized PGC2 loci genes over-represented in modules of co-expression by the modules’ relationship with polygenic risk score and diagnosis of schizophrenia. This additional step adds confidence that modules so identified are not likely to be based solely on illness state phenomena, many of which (e.g. treatment, chronicity effects) may be epiphenomena.

### Tier 1 genes

We identified 36 PGC locus genes that were distributed in one module (brown), significant for both diagnosis and association with genomic risk, i.e. PRS (**figure 3**). In principle, these genes and this network should bear an especially close relationship to schizophrenia pathogenesis and pathobiology. These genes are members of putatively relevant signaling pathways, such as PIK3/AKT signaling, which has numerous functions in neurodevelopment and adult brain and has been implicated in a variety of neurological and mental disorders, including SCZ [34]; a Ca^2+^ signaling pathway with numerous functions, including energetic metabolism [32] that was underscored in our study mainly by ATP2A2; a RAS/ERK signaling pathway represented by RRAS (RAS related small GTPase) implicated in cell adhesion and axon guidance [32].

Interestingly, when we interrogated STRING, a database of known and predicted protein-protein interaction [35] we noticed that several of the tier 1 genes are co-expressed in the same module with some of their interactors. For example, the PGC2 locus gene AKT3 is co-expressed with at least eight of its predicted interactors (ADCY2, CREB3L4, EIF4EBP1, GNB5, GSK3B, PHF20, PHLPP2, PIK3R1) [32, 35], and with serotoninergic receptors modulated by GSK3B (i.e. HTR2A). Another PGC2 locus gene, ATP2A2 (ATPase Sarcoplasmic/ Endoplasmic Reticulum Ca^2+^ Transporting), and its interactors (CALM1, RYR2, ITPR1) indicate potential dysfunctions on the Ca^2+^ Signaling Pathway in relation to energetic metabolism [32, 35]. NLGN4X (Neuroligin 4 X linked) previously implicated in autism and some of the interactors also members of the brown module (NRXN3, DLGAP1, DLG2, GRM1, GRM5) are constituents of the post-synaptic density and regulators of glutamatergic signaling [32, 35]. We highlight also RIMS1, co-expressed in brown module with RIMBP2 (RIMS Binding Protein 2) and synaptotagmins (SYT10, SYT11, SYT13, SYT16) [32, 35]. Some of the tier 1 genes in the brown module are in the RAS/ERK signaling pathway: RRAS (RAS related small GTPase) implicated in cell adhesion and axon guidance together with its interactors [35] and co-expression partners, i.e. PAK3 (role in dendritic development and synaptic plasticity), BRAF, RASGRP1, RASSF5, PRKCB (important for GABA-ergic synapse), RASAL2, RAPGEF2 (involved in neuritogenesis, neuronal migration), and RASGRF2.

The presence of intra-modular sub-clusters from different signaling pathways in our co-expression networks may indicate higher-order inter-network interactions. This scenario is plausible given previous studies which showed cross-talk between PIK3/AKT and RAS/ERK signaling pathways that regulates neurodevelopmental processes and synaptic plasticity [34]. This expected cross-talk between signaling pathways has been highlighted in a recent proposal about the underlying polygenic architecture of complex clinical syndromes [36].

### Tier 2 genes

We identified 53 potentially tier 2 PGC2 loci genes that were distributed in modules significant for diagnosis only, turquoise and yellow. A notable tier 2 gene in the turquoise module is ZNF804A, the first gene implicated using a GWAS approach to SCZ [33]. However, the mechanism by which ZNF804A is implicated in SCZ etiopathogenesis has yet to be determined, and earlier reports suggest association with a novel isoform only during fetal life [37]. Recent studies support the hypothesis that ZNF804A has multiple and important roles in neuronal physiology, including transcription regulation of interacting genes involved in cell adhesion, neurite outgrowth, and synapse formation [33]. Interestingly, we found that several genes which demonstrated transcriptional variation in studies based on ZNF804A knockdown were ZNF804A partners of co-expression in the turquoise module: C2Orf80 (unknown function), EIF4A2 (Eukariotic Translation Initiation Factor), and ATP1B1 (ATPase Na^+^/K^+^ responsible for maintaining the Na-K gradients across plasma membrane) [38, 39]. It is also interesting to note that in this large module, ZNF804A is just one of the 42 co-expressed transcription factors from the ZNF family. This implicates the formidable transcriptional regulation machinery that is putatively mobilized during various neuronal functions.

It seems worthy of comment that several historic candidate genes for SCZ are members of the modules containing tier 1 and tier 2 genes. Most notable examples are dopaminergic receptors (DRD4), receptor tyrosine kinases (i.e. ERBB4, receptor for neuregulins), NRG3 (growth factor that mediates cell-cell signaling and has multiple roles in neurodevelopment, and has been previously associated with SCZ [40], GABA receptors and glutamate decarboxylases involved in GABA synthesis (GAD1, GAD2), glutamatergic receptors, ionotropic and metabotropic (GRIA2, 3, 4, GRM1, GRM5, GRM8), serotoninergic receptors (i.e. HTR2A), COMT, and RGS4. The inconsistent results of previous studies on some of these genes, coupled with the lack of confirmation by GWAS, prompted a rebuke of the role played by historic candidate genes in the genetic risk for SCZ [41]. However, our results based on DLPFC co-transcriptome architecture suggest a complex distribution of genetic risk possibly organized in a modular fashion and indicative of vast gene-gene interactions that entrain new and previous genes associated with SCZ.

### Tier 3 PGC2 genes

We found that 59/148 PGC2 genes were enriched in two modules (blue and green) not related to the diagnosis of SCZ or to PRS. One of the stand-out tier 3 genes in the black module is CACNA1C, strongly associated with the risk for SCZ and bipolar disorder [42]. Other notable tier 3 PGC2 genes are CHRM4 (a drug target for SCZ [43], FURIN and TSNARE1. The latter two, FURIN and TSNARE1 were recently highlighted by Fromer et al [14] who showed in an experimental model of zebrafish neurodevelopment that overexpression of TSNARE1 and suppression of FURIN were associated with decrease of head size.

### Comparisons with previous work

Notwithstanding methodological differences, we found that our cortical gene expression networks were roughly consistent with previous similar studies. Of note, modules in our CAUC (CTRL+SCZ) network were significantly overlapping with modules originally reported by Oldham et al [7] (**figure 2**). Likewise, our results were in reasonable agreement with Fromer et al [14], considering the methodological differences related to mRNA processing, network construction and demographic characteristics of the samples. Interestingly, we found that 19 of the 31 PGC2 genes over-represented in their module of interest- M2C- were members of at least one of our tier genes. Moreover, nine of these 19 genes were actually tier 1 PGC2 genes in our study (**figure 3D**): SBNO1, TCF20, KCNB1, GRIA1, ATP2A2, HCN1, CSMD1, GRIN2A and NLGN4X. We also found a significant overlap between the modules of co-expression recently reported by Gandal et al (2018) [13] and our modules of co-expression (**supplementary table 6**).

### Methodological Considerations

Although these results are intriguing in many respects, we cannot rule out the possibility that they represent at least in part coincidental events, spurious associations, or effects of ongoing epiphenomena. Some of the limitations of our study are related to the network construction based on expression data at gene-level, which most probably obscures even more complex correlation patterns at the transcript and isoform level; likewise, we used RNA extracted from a tissue with a heterogeneous cell composition, which may not capture the cell-type specific co-expression architectures. We also cannot rule out the role of treatment exposure of SCZ samples or other epiphenomena in module construction in contrast to primary illness mechanisms. For example, animal studies have indicated some overlap between haloperidol regulation and co-expression networks enriched for SCZ genetic signals [12]. Further, while we have endeavored to pay special attention to the role of RNA quality as a confounder in co-expression, we cannot rule out this factor, and similarities to earlier work do not exclude a shared artifact. We have stressed genes in Tier 1 because of the convergence of association with illness state and also genetic risk, the latter not likely related to potential epiphenomena and confounders. In principle, genetic risk association obviates state only factors, but this is still conjecture. While modules of gene co-expression represent potential insights toward understanding physiological and etiopathogenic mechanisms, firm evidence of functional relevance requires experimental studies. Of note, our approach to network construction, based on enhanced adjustment for RNA quality, yielded a greater percentage of genes in grey, i.e. not in an explicit module, then in earlier studies. We believe this reflects several factors, including our relatively small sample size and removal of more complete co-expression based on co-degradation.

In conclusion, our study offers an extensive characterization of the co-transcriptome in the postmortem DLPFC of non-affected individuals and individuals with SCZ. Our results indicate potentially broad interactions of PGC2 locus genes, which may represent the tip of an iceberg of multiple convergent signaling pathways associated with the genetic risk of SCZ revealed through the co-transcriptome architecture in DLPFC. Interestingly, altered mechanisms suggested by these pathways span from prenatal neurodevelopmental events through brain functionality in adult life and hint not only to genetic factors, but also to an environmental contribution. Most importantly, our study highlights complex relationships between gene co-expression networks in the brain and the polygenic risk for SCZ and provides a strategy for using this information in selecting potentially targetable gene sets for therapeutic drug development.

## Acknowledgements

We are grateful for the generosity of the Lieber and Maltz families in establishing an institute dedicated to understanding the basis of developmental brain disorders. We would like to gratefully acknowledge the families of the subjects whose donations made this research possible. This research was funded by the Lieber Institute for Brain Development.

## Conflict of interest

The authors declare no conflict of interest.

---

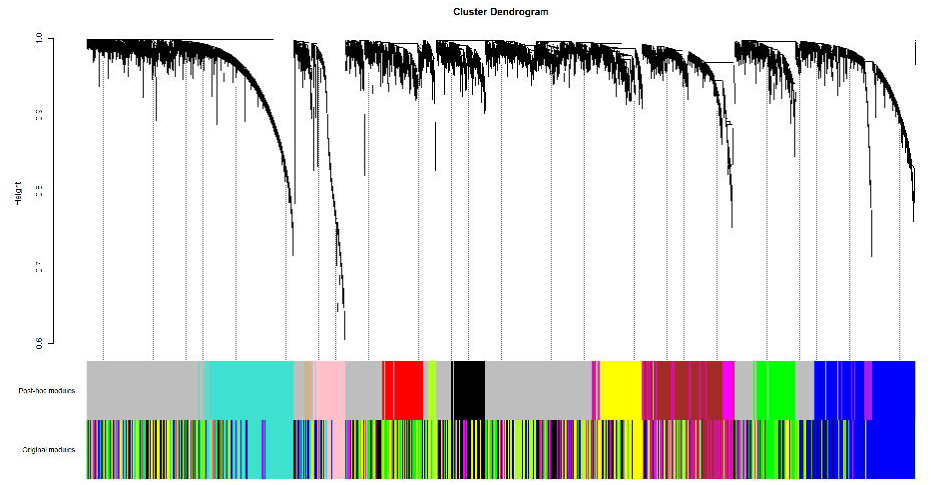

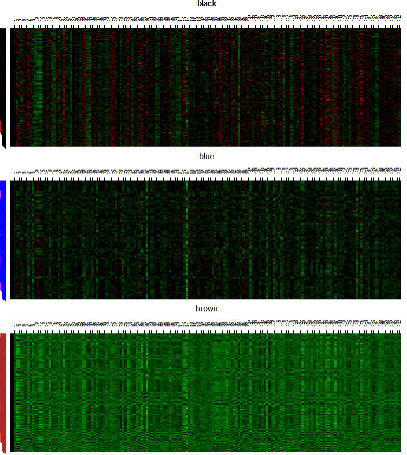

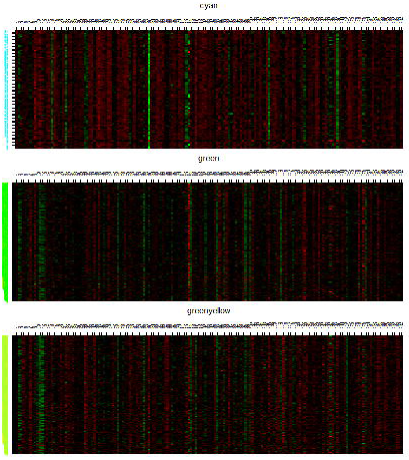

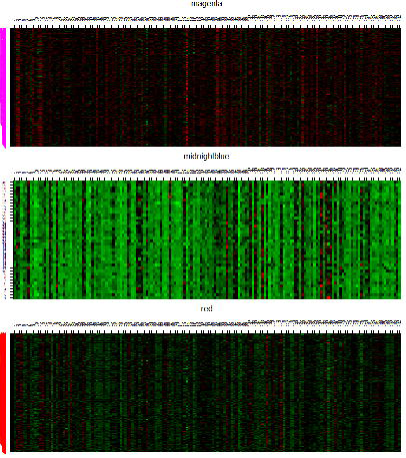

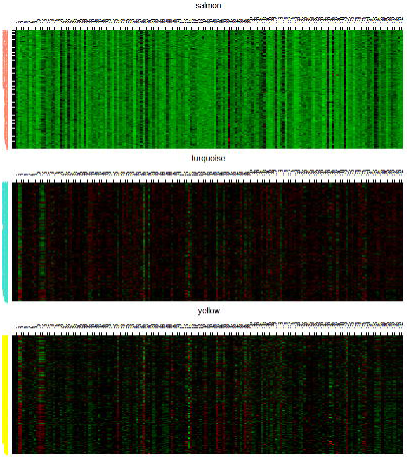

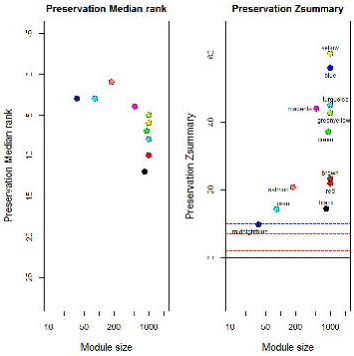

