## Supplementary Materials for "Identification and prioritization of gene sets associated with schizophrenia risk by co-expression network analysis in human brain"

**SUPPLEMENTARY METHODS**

Removing unwanted variance with Empirical Bayes linear models:

This method removes variation due to unwanted covariates, while preserving variance of covariates of interest. It uses Empirical bayes-moderated linear regression in a robust version, resistant to outliers of expression. However, it may be sensitive to outliers for the removed covariates; therefore we excluded the extreme outlying samples with respect to the removed covariates, which included RIN, postmortem interval (PMI), exonic mapping rate, alignment rate and 10 genomic principal components- snpPCs. For this analysis, no covariate of interest was “retained”, as we were interested only in removing potential RNA-quality artifacts.

Primary network construction: WGCNA steps

1. Adjusted expression data were used as input for network construction. Before selecting the beta power, the outliers for expression were detected by a fast algorithm of hierarchical clustering with “average” method.

2. Selecting the **beta power** by analysis of scale free topology for multiple soft thresholding powers.

3. Automatically creating the networks with *blockwiseModules* function. Parameters: correlation type = **bi-weight midcorrelation**; type of Toplogical Overlap Matrix (TOM) = **signed**; power=12 selected with soft thresholding to correspond to an **R^2≥0.8**; merging the modules whose eigengenes are highly correlated for a default height of 0.25. Minimum module size=40.

4. Assigning colors to module labels and saving the network output.

5. Additional checking the modules to see if eigengenes are correlated and merging modules that are too close.

Internal validation: Post-hoc co-expression network analysis controlling for “known” and “unknown” sources of variance

Controlling for “unknown” source of variance modelled by surrogate variables remain a work in progress in the context of co-expression network analysis (reference (14) in the main text). Nevertheless, because co-expression could reflect co-degradation, we applied a recently described approach based on an RNA degradation experiment to adjust for latent RNA quality not accounted for with observable metrics, such as RIN, mapping rate, mito-mapping, etc. (25 in main text).

In this study, we used a post-hoc network construction restricted to the genes clustered in modules of interest in the primary analysis, respectively 9239 genes that formed modules black, blue, brown, green, magenta, red, salmon, turquoise and yellow. Expression data for these filtered genes was adjusted with empirical Bayes linear modeling controlling for the same variables described in the main text (RIN, postmortem interval (PMI), exonic mapping rate, alignment rate and 10 genomic principal components- snpPCs) plus quality surrogate variables (qSVs) computed from a RNA degradation matrix as described in reference (25) in the main text.

The degradation matrix consisted of expression measures for 1000 chromosomal regions most susceptible to RNA decay when brain tissue from 5 donors was exposed to the room temperature for various intervals of time (25) in main text). Quality-surrogate variables computed from the degradation-matrix (“degradation principal components”- **qSVs** hence fort) were derived with the sva package in R, which implements a principal component analysis algorithm.

The number of qSVs was pre-determined with the num.sv function that uses the method of Buja and Eyuboglu (Buja,and Eyuboglu 1992 Multivariate Behav. 27:509-540) (option method=”be” in the num.sv() function). The new further “cleaned” data were used again in the *blockwiseModules* WGCNA function, with similar parameters like in the primary network construction, except for the power, which was set for a very lenient threshold of 5. The reason was that these genes were already known as being highly correlated and previously assigned in modules, and the sole purpose of analysis was to determine if the modular pattern of network is preserved in the context of a very conservative method of removing unwanted variance in RNA quality.

External validation: module preservation analysis

*ModulePreservation* function in WGCNA computes network-based, pair-wise module preservation statistics by taking as input adjacency matrices in a reference set and a test set. Most relevant output from the network preservation statistics is represented by measures of preservation for density (i.e. if a module remains densely connected in the test network and connectivity (i.e. if the hub gene status is preserved between reference and test networks, where hub genes are the most connected genes in a module). Four density and four connectivity preservation statistics are calculated in the module preservation analysis. Thresholds of preservation are determined by permutation tests. For this calculation, a default number of permutations was used (n=200). The output of this analysis reports observed and permutation Z score for each preserved measure in order to quantify the significance of preservation. Finally, the individual Z scores are summarized in a composite measure called Z_summary_. By convention, Z_summary_ indicates no preservation for a range between 0-2, weak preservation between 2-10 and strong preservation >10. Another composite statistics is the median rank, which is based on the rank of observed preservation statistics and consequently it does not require a permutation test. Likewise, in contrast to the Z_summary_, it is not dependent on the module size (the number of genes in a module). However, the disadvantage is that it is applicable only to the ranking modules. Therefore, both measures- Zsummary and median rank- give important information about the preservation of modules from the reference network in the test network. Details about the computation of preservation measures are presented in (24).
